# Propagation dynamics of electrotactic motility in large epithelial cell sheets

**DOI:** 10.1101/2022.07.30.502112

**Authors:** Yan Zhang, Guoqing Xu, Jiandong Wu, Rachel M Lee, Zijie Zhu, Yaohui Sun, Kan Zhu, Wolfgang Losert, Simon Liao, Gong Zhang, Tingrui Pan, Zhengping Xu, Francis Lin, Min Zhao

**Affiliations:** Department of Ophthalmology & Vision Science, University of California, Davis, CA 95616, USA.; School of Public health, Hangzhou Normal University, Hangzhou 310018, China.; Institute of Environmental Medicine, Zhejiang University School of Medicine, Hangzhou 310058, China.; Micro-Nano Innovations (MiNI) Laboratory, Department of Biomedical Engineering, University of California, Davis, CA 95616, USA.; Department of Physics and Astronomy, University of Manitoba, Winnipeg, MB, R3T 2N2, Canada.; Department of Applied Computer Science, University of Winnipeg, Winnipeg, MB, R3B 2E9, Canada.; Institute for Physical Science and Technology, University of Maryland, College Park, MD 20742, USA.; Department of Physics, University of Maryland, College Park, MD 20742, USA.; Brain Engineering Center, Anhui University, Hefei 230601, China.; Institute of Biomedical and Health Engineering, Shenzhen Institute of Advanced Technology, Shenzhen 518055, China.; Shenzhen Engineering Laboratory of Single-molecule Detection and Instrument Development, Shenzhen, Guangdong 518055, China.; Suzhou Institute for Advanced Research, University of Science and Technology of China, Suzhou 215123, China.; Department of Precision Machinery and Precision Instrumentation, University of Science and Technology of China, Hefei 230026, China.; Department of Dermatology, University of California, Davis, CA 95616, USA.; Lead contact.

**Keywords:** epithelial sheet, collective cell migration, galvanotaxis, electrotaxis, cell motility, cellular interplay, cell junction, wound healing, keratinocytes, human

## Abstract

Directional migration initiated at the wound edge leads epithelial sheets to migrate in wound healing. How such coherent migration is achieved is not well understood. Here we used electric fields to induce robust migration of sheets of human keratinocytes and developed an *in silico* model to characterize initiation and propagation of epithelial collective migration. Electric fields initiate increase in migrations directionality and speed at the leading edge. The increases propagate across the epithelial sheets, resulting in directional migration of cell sheets as coherent units. Both the experimental and in silico models demonstrated vector-like integration of the electric and default directional cues at the free edge in space and time. The resultant collective migration is remarkably consistent in experiments and modeling, both qualitatively and quantitatively. The keratinocyte model thus faithfully reflects key features of epithelial migration as a coherent tissue *in vivo*, e.g. that leading cells lead, and that epithelium maintains cell-cell junction.

## Introduction

Large epithelial sheets migrate directionally into wounds, which is a hallmark of wound healing (Eming et al., 2014; Friedl and Gilmour, 2009; Gurtner et al., 2008; Ilina and Friedl, 2009; Martin, 1997; Pastar et al., 2014; Rorth, 2012; Zhao et al., 2003). In this type of collective migration, wound edge cells lead; cells maintain their relative positions and intercellular junctions; and cells migrate as a continuous sheet, thus there is integrity of the epithelium. In corneal wound healing, for example, wound edge cells lead migration into the wound, and follower cells migrate in a highly coherent manner with less than 3% of cells changing their relative positions in epithelial sheets (Safferling et al., 2013; Zhao et al., 2003). Such migration can be observed at least millimeters away from the wound edge and is essential for wound healing.

Electric fields (EFs) are found in wounds. Numerous experiments over nearly two centuries from different laboratories have demonstrated the existence of the wound EFs (Barker et al., 1982; Foulds and Barker, 1983; McCaig et al., 2005; Mukerjee et al., 2006; Nuccitelli et al., 2008; Reid and Zhao, 2011). Remarkably, sheets of corneal epithelial cells respond to applied EFs by directional collective migration (Zhao et al., 1996; Zhao et al., 2006). Such collective migration has been elegantly demonstrated in other types of cells with advanced engineering devices (Cohen et al., 2014; Shim et al., 2021; Zajdel et al., 2021; Zajdel et al., 2020). Perhaps due to different cell types or culture conditions, very interesting cell behaviors have been demonstrated in collective electrotaxis. For example, leading-edge cells of elegantly engineered MDCK cell monolayers were insensitive to the applied EFs and thus migration was largely limited to the inner region cells without net displacement of the entire cell sheet (Cohen et al., 2014). In primary cultures of mouse keratinocytes, EFs resulted in catastrophic damage (cell death) to the leading edge (Shim et al., 2021). In some models of the collective electrotaxis, cells do not maintain monolayer property, i.e. cell-cell junctions are not maintained (Zajdel et al., 2020). Those cell behaviors are in contrast to in-vivo observations where wound edge cells lead and whole-cell sheets migrate directionally (Eming et al., 2014; Gurtner et al., 2008; Martin, 1997; Pastar et al., 2014).

Directional migration initiated at the wound edge leads collective migration of epithelial sheets to migrate into the wounds thus is critical for wound healing. How such coherent migration across large space (mm) and time (hours) is achieved remains not well understood. We, therefore, sought to develop an experimental model of collective electrotaxis that share those important features in collective migration of epithelial sheets in wound healing. A free edge, which is usually produced by scratch wounding or removing barrier next to a confluent monolayer, is a combination of multiple guidance cues, including space availability, population pressure, contact inhibition release, and free edge dependent activation of EGFR (Block et al., 2010; Klarlund and Block, 2011). Compared with other direction signal, default directional cues at the free edge has been shown in many types of cells, continuously present, and easy to be engineered to induce uniform directional cell migration. We used engineered monolayers of human keratinocytes (HaCat cell line) with well patterned free edges and applied EFs of physiological strength to induce collective electrotaxis of sheets of human keratinocytes. In parallel, we developed a mathematical model to capture and predict the key parameters that determine collective cell behaviors. Results from experimental and *in silico* models, in agreement qualitatively and quantitatively, reveal EF-induced initiation of directional migration at the leading edge, which then propagate across the epithelial sheets that maintain cell-cell junction, resulting in coherent collective migration of the whole-cell sheets. Both models demonstrate vector-like integration of electric guidance and default directional cues at the free edge in space and time, with an electrotactic delay effect.

## Results

### Robust electrotaxis of epithelial sheets of human keratinocytes

To establish a migration model of epithelial sheets *en masse* with intercellular junction and geometry integrity, we cultured human keratinocytes (HaCaT) into epithelial sheets of various shapes and sizes using a microfabricated PDMS stencil with the desired shape, for example, a square shape of up to 2 mm × 2 mm size (Fig. 1A (1 mm × 1 mm), Fig. S1A (2 mm × 2 mm)). Cell sheets of rectangular, circular, triangular, and other shapes and sizes were also made as needed (Fig. S1B). These cell sheets formed tight cell-cell contacts typical of a monolayer epithelium. Upon stencil removal, cells at the edge followed the default directional cues at the free edge to migrate into the acellular area with a migration direction perpendicular to the free edge (Fig. 1A, B; Video S1).

**Fig. 1.**
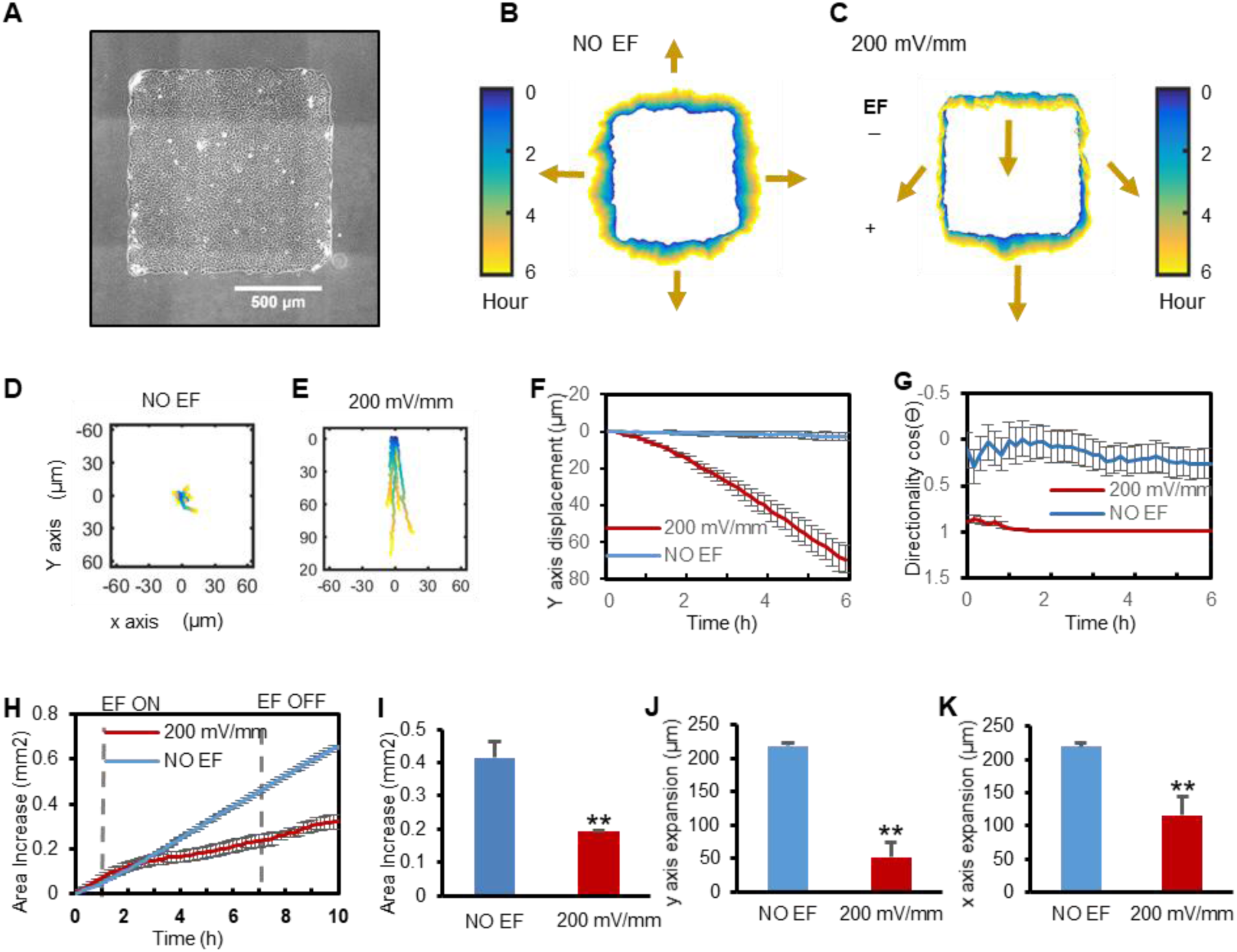
EFs induce collective electrotaxis of large epithelial sheets that maintain cell-cell junctions. **A** Phase contrast image of square cell sheet. Scale bar = 500 µm. **B-C** Collective electrotaxis demonstrated by contour assay showing color-coded edge position of cell sheets. Color coding from blue (0 h) to yellow (6 h). Arrows indicate overall edge migration direction and distance. An EF guided directional migration of the cell sheet downward (**C**), whereas a control cell sheet without EF expanded in all direction (**B**). Polarity and strength of the field as shown. **D-G** Collective electrotaxis of whole cell sheets shown by the trajectories of the geometric center (centroid) of cell sheets (n=6) (**D**, **E**). Y axis displacement (**f**), directionality along the field line (**G**), as a function of time. Directionality is calculated as cosθ, where θ is the angle between each centroid trajectory and the electric field line. Data are mean ± SEM from 6 independent experiments. Positive value of cosθ indicates migration to the anode; negative to the cathode, zero indicates random direction. **H-K** EFs maintained cell sheet geometry by suppressing dispersal of cells. Quantification of area increase (**H**-**I**), y axis expansion (**j**), and x axis expansion (**K**). (\*\**p*<0.01). Data are mean ± SEM from 6 independent experiments. See also Figure S1, Video S1.

When an EF was applied to the cell sheets, migration *en masse* became evident within 30-60min, with the leading edge (the free edge with default migration direction to the anode) migrating towards the anode, the rear edge (the free edge with default migration direction opposite to the cathode) retracting in order to migrate towards the anode, and the whole sheet migrating directionally (Fig. 1C; Video S1). The geometric center of the cell sheets moved directionally (∼80 µm) in an EF (Fig. 1E) in contrast to no net displacement (∼0 µm) in the no EF control (Fig. 1D). A drastic difference in the movement of the geometric center of the cell sheets was clearly induced by the applied EFs (Fig. 1F, G).

To examine the integrity of the cell sheets during migration *en masse* in EFs, we recorded the leading region, center, trailing region of the cell sheets at high magnification. Cells maintained tight cell-cell contacts with less than 10% of cells changing their relative position and no instances were observed of gap spaces appearing between cells (Video S2). Cell sheet contours revealed that the shape and size of cell sheets were maintained significantly better in EFs than those cell sheets without exposure to EFs (Fig. 1B, C, H-K). The cell sheets thus maintain monolayer integrity in EF-guided migration *en masse*. This is particularly important because when epithelial tissues move to heal a wound, integrity (barrier function) should not be compromised.

### Leading edge cells lead the whole cell sheet migration in EFs

To determine the time course of cell response in epithelial sheets, we analyzed migration trajectories of individual cells in four regions (Rear region, Central region, Leading region, Right/Left region) with higher magnification videos (Fig. 2A; Video S2). Leading region cells showed a directional migration to the guidance of EFs as its original migration direction agree with the EF guidance, while cells in other regions take time and gradually assume the direction to migrate in the same direction as the leading edge cells. (Fig. 2B-E). These regions showed a distinct migration trend compared with the no EF control cell sheet (Fig. 2F). Cells in the leading region migrated towards the cathode first while rear region cells followed with a lag of ∼20 min in directionality and speed in the cell sheet (Fig. 2B-D). Cells in the center region of cell sheets notably had almost the same magnitude of Y axis displacement towards the anode as the leading region cells (Fig. 2B). Cells in the side region migrated with a consistent bias along the EF lines (Fig. 2E).

**Fig. 2.**
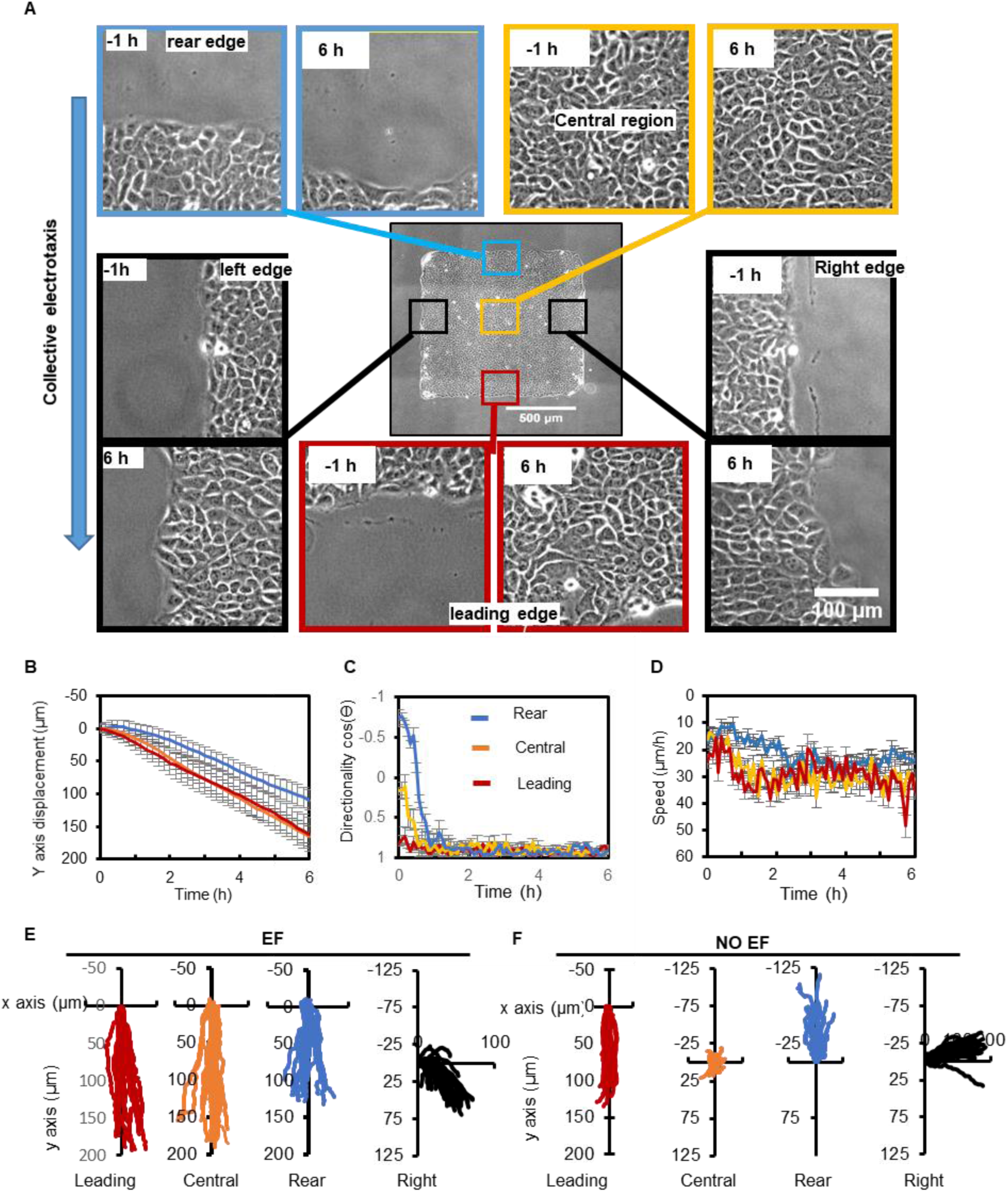
Leading cells lead collective electrotaxis with suppressed migration sideways. **A** Snapshots of regions of a cell sheet 1 hour before the onset of EF and 6 hours in the EFs. **B-D** Cells in the leading (L), Central (C), Rear (R) regions were tracked, for Y axis displacement (**B**), directionality (**C**), and speed (**D**) as a function of time during EF application. Data are presented as mean ± SEM from 20 cell migration trajectories from three independent experiments. **E-F** Migration trajectories of cells in leading region, central region, right region, and rear region of the cell sheet in an EF and NO EF group. Data from a representative of three independent experiments are represented (n = 20 for each group). EF = 200 mV/mm. See also Figure S7, Video S2.

### EFs induce a wave-like propagation pattern of directionality and speed across the cell sheet

To understand how cell sheets respond to electrical guidance, we utilized particle image velocimetry (PIV), which provides a quantitative visualization of directionality and speed of the movement of whole cell sheet (Fig. S2). PIV revealed that the cell sheet exhibited an initial high motility ring at the edge of the cell sheet followed by a motility wave during the subsequent outward expansion: When there were no EFs, the directionality of cell movement into the cell-free area first increased at the edge of the cell sheet and then propagated into neighboring cells (Fig. 3A, Video S3). To better characterize the systematic evolution of motility patterns in the cell sheet, we averaged those variables over the observed cells in the x axis, thereby reducing the dimensionality of the system to only one spatial dimension and one temporal dimension (see Methods). These kymographs movie vividly showed the directionality cue transition: two directionality waves developed at the edges of the cell sheet, then propagated towards the center of the cell sheet, and finally rebounded back when two waves met (Fig. 3C). When the cell sheet was exposed to an EF of 200 mV/mm, these directionality waves were disturbed in a polarized fashion. The directionality waves were inhibited at the rear but persisted at the front of the cell sheet. The EF-induced directionality wave initiated at the leading edge of the cell sheet and then propagated to the rear edge of the cell sheet; the duration of the propagation was about 20 minutes (Fig. 3B-C, Video S3). Interestingly, the directionality waves coming from the edges perpendicular to the EFs were inhibited, limiting side edge expansion (Fig. S3A). Cell migration speed accelerated upon electrical stimulation (Fig. 3B, 3D, Fig.S3B, Video S4).

**Fig. 3.**
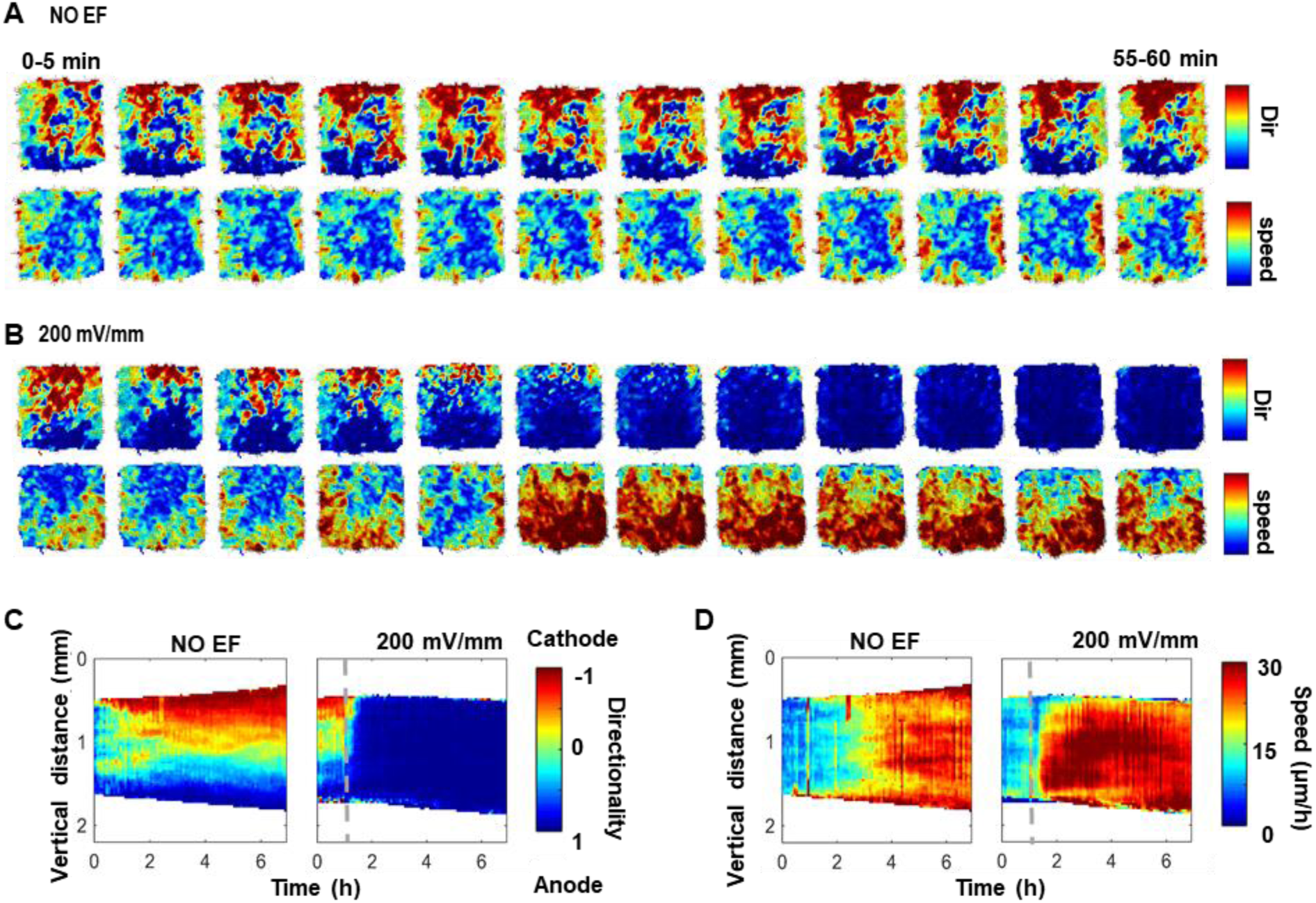
EFs, overriding default directional cues at the free edge, guide and mobilize cell sheets by inducing propagation of increased directionality and speed across the whole cell sheets. A-B. Snap shots of directionality (Dir) and speed heatmaps of a cell sheet in the first hour after EF application. The images are derived from **Video S3** and **Video S4**. Directionality and speed of movement of the cell sheet shown as heatmaps from PIV analysis from two adjacent images with a time interval of 5 min. **C-D** kymographs of directionality and speed as a function of time along the x axis (**C**) and y axis (**D**), respectively. Kymographs in no EF group show that directionality and speed propagate from the free edge to the center of the cell sheet and then deflecting back when they meet. Kymographs in an EF (200 mV/mm) show a dramatic increase in directionality and speed propagating from the leading edge to rear edge. Dashed lines indicate the onset of the field. The experiment duration is 7 hours. Kymographs presented are drawn from one experiment and confirmed in three independent experiments. See also Figure S2, S3, Video S3, S4.

### A PBC (Particle-Based Compass) model successfully reproduces the propagation dynamics of the electrotactic motility wave

In our previous research, we demonstrated that a simple particle-based compass (PBC) model was sufficient to successfully capture motility waves during monolayer wound healing (Zhang et al., 2017). Briefly, cells in the model are represented as particles. Cellular interplay was based on cell-cell distance such that varying the cell-cell distance could result in repulsive or attractive interactions. A free edge signal representing default directional cues at the free edge was presented to bias cell migration toward the free space. Cell migration speed was calculated independently from cell migration direction. Cell migration direction was determined based on the sum of the neighboring cell-cell interactions (Fig.4C). In this previous model, EF guidance was not considered.

**Fig. 4.**
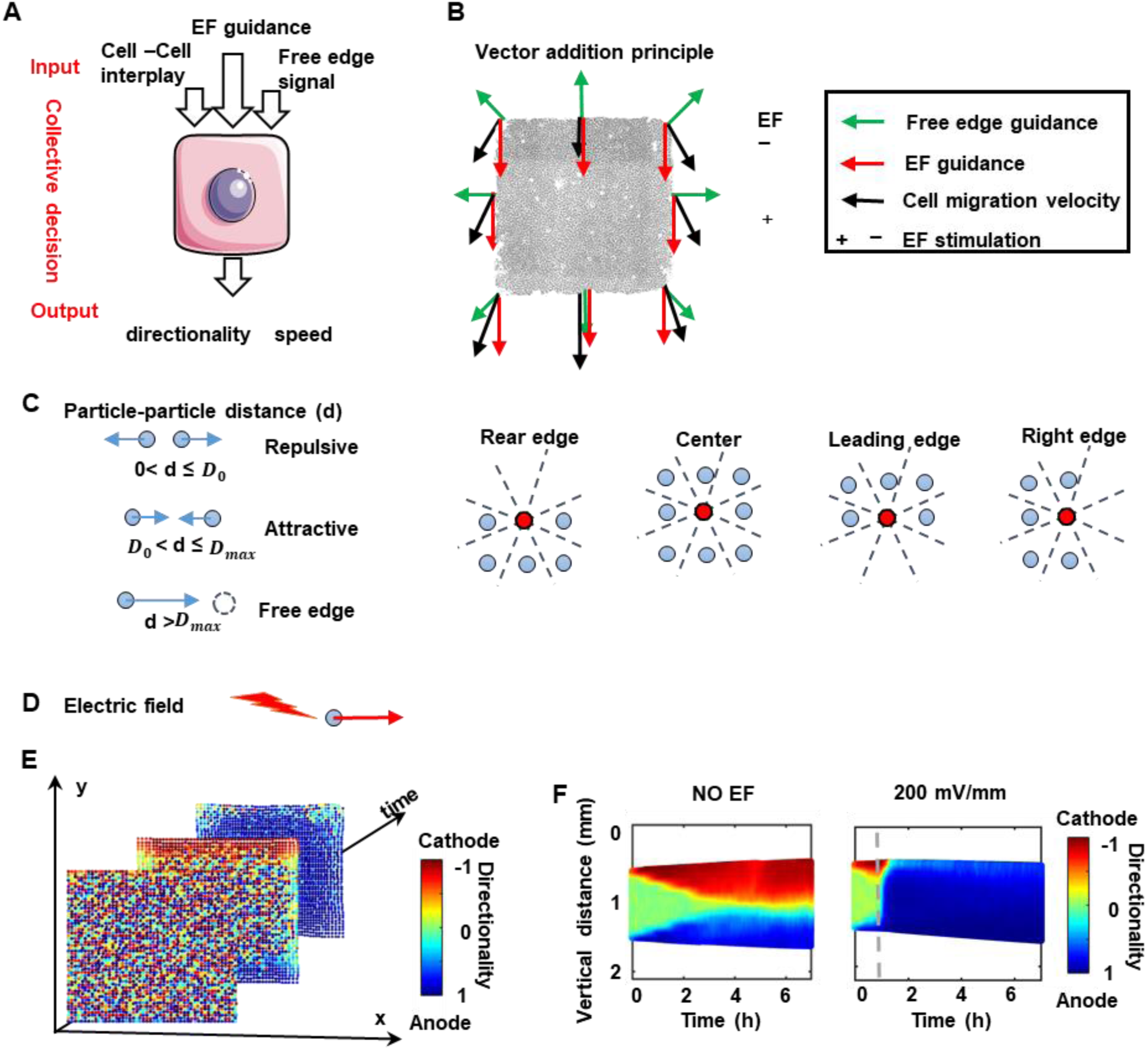
*In silico* model captures the propagation dynamics of collective migration. **A, B** Directional cues (EF, free edge, and cell-cell interplay) as inputs, and speed and directionality as outputs. Vector addition as a principle in migration directionality determination. **C** A elastic ball representing a cell. The cell–cell distance *d* determines the cellular interaction, where *D_0_* and *D_max_* denote the threshold of repulsive interactions and the threshold of the free edge effect. Red balls at the rear edge, leading edge and right edge have three vacant neighbor sectors out of eight, which bias the direction of migration. **D** Directional guidance of EF on the particle. An EF biases the directionality. **E** A set of snapshots of the computer simulation of collective cell migration of a cell sheet (time=0 h, 1 h, 2 h) of 2500 cells. The snapshots are derived from **Video S5**. **F** *In silico* kymographs of cell migration directionality for NO EF and 200 mV/mm EF groups. Color codes the directionality as shown, compare with Fig. 3C. See also Table S1, Figure S4, Video S5.

In the current study, we patterned cell sheets into a well-defined square shape and found distinct responses of cells in different regions of the cell sheet, coherent directional migration of the whole cell sheet in response to EFs, and a wave-like propagation pattern of directionality and speed. Thus, we hypothesize that cells in the cell sheet integrate multiple directional signals by following the vector-like addition interaction.

To test this hypothesis, we developed the previous PBC model into a two-dimensional model and took into account EF guidance (see Method and Supplementary Material for details). In this model, each cell can process the given inputs of EF guidance, default directional cues at the free edge, and cell-cell interplay to generate outputs such as speed and directionality based on a set of defined rules (Fig. 4A-B). The model treats the cell sheet as an array of elastic balls (Fig. 4C-E). All cells in the cell sheet equally sense the EF signal. This is a common feature that cells often respond to changes of external signals in a time-delayed manner. To model the delay effect, the model cells do not respond immediately to changes in the guiding signals. Instead, a delay variable is defined to allow the cell to retain a certain portion of its previous migration speed and directionality. On the other hand, when the EF is turn on or turn off, the strength of EF increases or decrease linearly in our model. The updated migration state is calculated based on the 2D force vector-like interaction. To simplify the model, we set cell speed to be a constant plus noise in the model. Computer simulations of this model show similar patterns of directional collective migration as our cell culture experiments. Cells close to the free edge migrate directionally first and the cells behind follow in a time-dependent manner (Video S5). Kymographs of simulated directionality showed wave patterns similar to the experimental data (Fig. 4F, Fig. S4). Thus, a 2D PBC model that vector integrates default directional cues at the free edge and EF cue in combination with cell-cell interplays can reproduce the experimentally observed directionality wave pattern, suggesting its potential as a working model for describing collective epithelial electrotaxis and making further predictions.

### The PBC model predicts a three-phase directionality pattern in cell sheets

We further tested the viability of the PBC model in predicting how changes in the EF affect the directionality patterns in cell sheets. When a lower EF combing with a longer response time (2 hour) was applied in the model, the simulation showed that a similar directionality wave developed from the leading edge but propagated much more slowly than in the high EF parameter condition, which is experimentally equivalent to 200mV/mm (Fig. 3C vs. Fig. 5A). Taking both the high EF and low EF groups into consideration, the wave-like propagation pattern can be roughly described by three phases that differ in wave propagation rate as the wave moves across the cell sheet: the initial phase (P1), ramping-up phase (P2), and saturation phase (P3) (Fig. 5A). In the initial phase, directionality begins to propagate away from the free edge (P1), followed by a faster phase of spreading throughout the cell sheet (P2). At longer times, during the saturation phase (P3), the directionality wave spreads to the rear edge of the cell sheet. Compared with the wave-like propagation pattern in the high EF (Fig. 3C), the duration of P1 and P3 in the low EF was extended. The extension of P3 is much more pronounced than the change in P1 (Fig. 5A). This prediction from the model was verified experimentally using a 50 mV/mm EF (Fig. 5B). Under 50 mV/mm EF stimulation, directionality towards the anode did not spread to the whole cell sheet within the first 6 hours of the experiments. Directionality within the cell sheet was largely hindered by the tug-of-war of direction signals at the rear edge of the cell sheet, as shown by a comparison to the low EF single cell data (Fig.5E) and as predicted by the PBC model with low EF parameters (Fig. 5A). Kymographs of both the simulations and the experiments consistently showed the cell sheet expanding in the direction perpendicular to the low EF (Fig. S5A-B, Video S6).

**Fig. 5.**
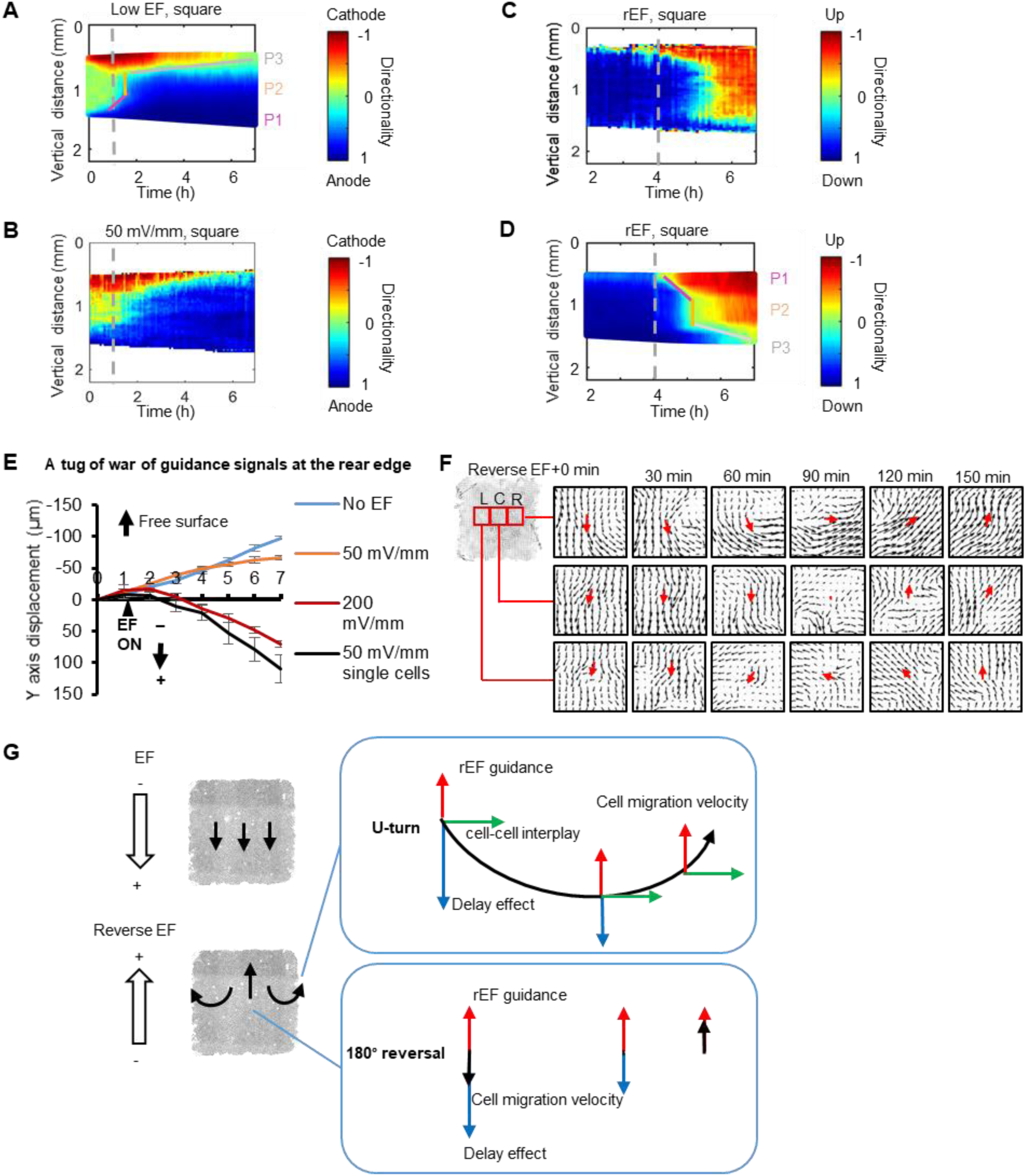
*In silico* model predicts and reproduces spatiotemporal dynamics of collective electrotaxis of cell sheet. **A-D** *In silico* spatiotemporal dynamics of collective electrotaxis (**A**, **D**) are in good consistency with experiments (**B**, **C**). For low EF group (**A**, **B**), the experiment includes 1 hour no EF, 6h for low EF. For reverse EF group (**D**, **E**), the EF polarity was reversed after the cell sheet assumed steady overall directional migration for 2 hours. Dashed lines indicate the onset (**A**, **B**) and reversal of the field polarity (**D**, **E**). Kymographs presented are representative of three independent experiments with the same results. Cell sheets of 2500 cells. The wave-like spatiotemporal dynamics of collective electrotaxis can be roughly described by three phases with distinct slopes: the initial phase (P1), ramping-up phase (P2), and saturation phase (P3). **E** A tug of war of the guidance signals. Y axis displacement of cells at rear edge of the cell sheet at 0, 50, 200 mV/mm are compared to that of cells in sparse culture in a field of 50 mV/mm. **F** Cells respond to reversal of EF by making region dependent turns: PIV of cell cluster at right (R), left (L) and central region(C). The direction of vector flows indicates the directions of cells migration. The length of the vector indicates the migration speed. Typical data from one of three independent experiments. Cells at right and left regions of the cell sheet make U-turns following EF reversal. Cells in the center of the cell sheet make sharp 180-degree reversal of migration direction. The red arrows indicate the main direction of the vector flows in the regions. **G** Hypothetical schematics show U-turn and 180-degree reversal following EF polarity switch. Blue arrows indicate momentum of default cell migration (delay effect, which gradually decays), red arrows new EF guidance, green arrows free-edge direction (cell-cell interplay), and black arrows cell migration velocity. See also Figure S5-6, Video S6-8.

Furthermore, we tested how parameters in the model affect the three-phase directionality pattern. First, we set the EFs at high EF level, tuned the cellular response time form 0 min to 120 min, and we found that total and duration of each phase of the directionality pattern were elongated (Fig.S6A). Next, we set the cellular response time to EFs to 2 hours, and varied the EFs from low EF to high EF. Simulation results indicated that the higher EF, the shorter duration of the three-phase directionality pattern, and that of third phase (Fig. S6B). Last, we tested whether the existing motility wave induced by free edge affected the three-phase directionality pattern. We started EF stimulation at 0, 1 and 2 hours and results suggested that the existing motility wave affected the third phase of the directionality pattern (Fig. S6C). Together, our PBC model indicates that the three phased directionality pattern induced by EFs are affected by the strength of EFs, cellular response time to EFs and motility state before EF stimulation, which are consistent with the experimental results.

### The PBC model reproduces the electrotactic delay of cell sheet qualitatively and quantitatively

We next tested the PBC model’s ability to capture the delayed effect of cell sheet electrotaxis in response to reversal of a high EF stimulus (experimentally equivalent to 200 mV/mm). The 7-hour experiments started with one hour without an EF, followed by 3 hours of a direct current EF and a subsequent 3 hours of a direct current EF in the opposite direction. For simplicity, kymographs present the data from the second hour to the end of the experiment. After 3 hours in a high EF, directionality of the cells across the whole cell sheet turned to the anode (blue), and after reversal of the EF direction, we observed that cells gradually turned to the new anode (red), and this trend started with the new leading edge (the previous rear edge). The directionality dynamics in the kymograph could be separated into the three phases in the same fashion as described above: initial phase (P1), ramping-up phase (P2), and saturation phase (P3), which show both spatial and temporal features (Fig. 5C, Fig. S5D). The new leading edge responded to the reversed EF first. Cells located at the side edge (parallel with the EF) also responded to the reversed EF earlier than the cells in the inner region and at the new rear edge of the cell sheet. The previous leading-edge cells continued migrating to the previous anode of the EF (Fig. 5C, Video S7). Kymographs perpendicular to the electric fields, unexpectedly revealed that, the expansion tend to begin at the edge parallel with EF line and propagate inward to the cell sheet but is mainly restricted to the region close to the free edge (Fig. S5D).

By gradually decreasing the strength of the EF to 0 and then increasing the strength of the EF in the opposite direction combing adjusting the repulsive threshold distance threshold from 0.1 to 0.13 after reversing EF in the model, we successfully reproduced the motility pattern agree with the in vitro cell experiments (Fig.5C vs. Fig.5D; Fig.S6C vs. Fig.S6D). Our model simulation suggests the role played by cell-cell interaction (repulsive threshold distance threshold in mathematical model) in mediating collective electrotaxis in response to reversed EF.

Snapshots of directionality heatmaps of cell experiments further revealed that when EF guidance reverses, cells located at different regions of cell sheet respond to this instantaneous change of direction cue differently. The majority of cells in the cell sheet maintained their original migration behavior for the first hour following EF reversal, suggesting a delayed effect of EFs on cell sheet migration. Regions in the center of the cell sheet reversed migration direction gradually after EF reversal, while regions at right and left edges the cell sheet gradually make U-turn after EF reversal (Fig. 5F, Video S8). Together with the tug of war we observed at the rear edge of the cell sheet (Fig. 5E) and the region dependent response to reversed EF (Fig. 5F), our experimental results further consolidated our hypothesis of the vector-like interaction of for cellular integration of multiple directional signals in collective electrotaxis. In this *in silico* model cells in sheet mainly respond to the reversed EF guidance, delay effect, and cell-cell interplay coming from the left and right edges. For cells at right region of the cell sheet, cells have cellular interplay guidance to migrate towards the right edge. As the delay effect decays gradually, cell migration velocity is overtaken by the reversed EF guidance and cell-cell interplay, and thus the region makes a U-turn. For cells in the central region of the cell sheet, the cell-cell interplays that come from the right and left edge are balanced and thus have a net zero effect. As a result, the cell migration velocities of the cells in the center of the cell sheet are determined by the reversed EF guidance and the delay effect. As the delay effect decays gradually, cell migration velocity is overtaken by the reversed EF guidance, and thus the region makes a 180-degree reversal (Fig. 5G). Thus, our PBC model successfully captured the delayed response to EF direction reversal.

### Convergent EFs induce converging collective electrotaxis

Endogenous electric fields at a wound all pointing to the wound center, and cells move to the wound center where electric fields “converge”. Those endogenous EFs and convergent cell movement are observed in skin wound healing in vitro (Barker et al., 1982; Nuccitelli et al., 2008; Reid et al., 2007; Reid et al., 2005). We therefore tested whether convergent collective electrotaxis could be induced by convergent EFs, i.e. establishing EFs all pointing to the center of a circle. A vacuum-assisted assembly method was utilized to allow a multi-electrodes microfluidic chip to direct attachment of a confluent cell monolayer and pattern the convergent EFs (Fig. 6A-C). The convergent EFs induced converging collective electrotaxis. In a three electrodes system, Line Integral Convolution (LIC) visualization of cell movement shows collective migration of cells followed very well the field lines that converge at the center (Fig. 6C-D).

**Fig. 6.**
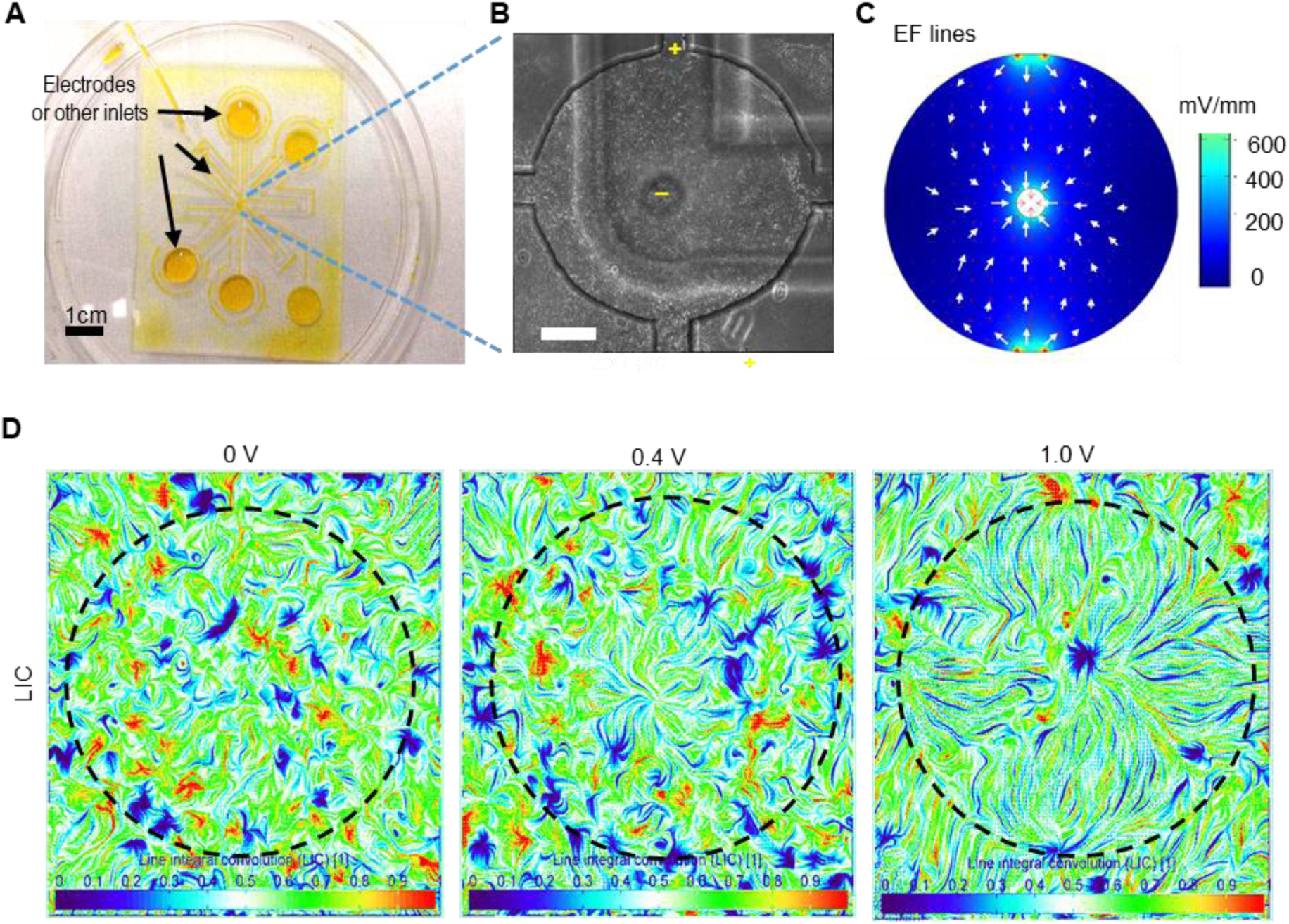
Convergent EFs induce converging collective electrotaxis. **A** Microfluidics design to establish convergent EFs. A multi-electrodes microfluidic chip was placed over a cell monolayer. In the view, there were five channels - four of them connected to the edge of the round platform, one of them suspended on the top of the platform. Channels were filled with cell culture medium and could be selectively connected to biocompatible electrodes. **B** A phase contrast image of cell monolayer with multi-electrodes shown. In this experiment, three of channels were selected. The one in the center of the platform served as the cathode, the other two served as anodes. Bar = 250 µm. **C** Convergent EF lines simulated by COMSOL Multiphysics. The intensity of EFs is in the range of 0∼400 mV/mm. **D** Representative Line integral convolution (LIC) images of collective electrotaxis in convergent EFs as in (**C**). The voltage potential between anode and cathode was 0 V, 0.4 V, 1.0 V, respectively. LIC is a technique to visualize vector fields, which show convergent migration development when the field strength increases. Color encodes the index of LIC.

## Discussion

### Collective electrotaxis of sheets of keratinocytes

Epithelial migration as a continuous sheet is important in wound healing and development. Several in vivo and in vitro models have been used to study collective migration of epithelial sheets. Robust collective electrotaxis of large sheets were demonstrated using corneal epithelial cells (Zhao et al., 1996; Zhao et al., 2006). Collective electrotaxis have been studied using MDCK cells and mouse primary keratinocytes with interesting features, such as leading edge cells being insensitive, no overall migration of the whole epithelial sheets, breaking down of cell-cell junction, and dying of leading edge cells in an applied EF (Cohen et al., 2014; Shim et al., 2021; Zajdel et al., 2020). Those are not observed in collective migration of epithelial sheets in various wound healing models in vitro and in vivo, nor in collective electrotaxis of corneal epithelial sheets (Ud-Din and Bayat, 2017; Zhao et al., 1996; Zhao et al., 2006). In this study, we used a well-established human keratinocyte line to generate large epithelial sheets that maintain cell-cell junction, have robust leading-edge migration and show no cell death in EFs to characterize the spatiotemporal dynamics of migration speed and directionality in electrotaxis. In parallel, we have developed an *in silico* model to determine key parameters in collective electrotaxis. The experiment model and *in silico* model produce results that match each other qualitatively and quantitatively. Results from the experimental and *in silico* models together reveal important spatiotemporal dynamics in collective electrotaxis, and critical mechanisms mediated by position of cells in related to other cells in the cell sheet, temporal information of cell migration speed and direction, cell-cell junction, and integration of different directional cues.

### Cell sheet integrates default directional cues at the free edge and electrical cue by vector-like superposition

It is not clear how cells integrate directional cues when making collective decisions, especially when EFs are present with other cues (Rodriguez and Schneider, 2013). There are several scenarios that could exist when cells integrate multiple directional cues. The first is a simple vector addition of the migration responses. When cues are aligned, cooperation occurs, and an additive effect on the cell migration speed and/or directionality is produced. When the angle is 180°, competition between cures occurs, while intermediate angles result in an intermediate scenario (Rodriguez and Schneider, 2013). Another theory for cue integration is hierarchical dominance. This was first found in the decision-making process for immunological chemotactic cues presented to neutrophils. Neutrophils migrated preferentially toward target chemoattractants rather than intermediary endogenous chemoattractants (Foxman et al., 1997; Heit et al., 2002; Lin et al., 2005).

Taking advantage of detailed spatiotemporal analysis of collective migration, we provide experimental evidence supported by *in silico* results that cells at the boundary of a large cell sheet are sensitive to EFs, initiate and lead collective electrotaxis. Those cells appear to employ a vector-like supposition when coupling free edge signals and electric guidance. The front edge of the cell sheet leads the whole cell sheet and undergoes more rapid and efficient directional migration in response to the applied EF compared to the rear of the cell sheet. At the rear of the cell sheet, there is a tug of war between the EFs and directional migration initiated by contact inhibition of locomotion (Fig. 5E). When EF lines are parallel with the free edge, cells bias to the guidance of electric cues depending on the field strength (Fig. 2E). For cells located in the cell sheet, cell responded to a reversed EF by making a U-turn or 180-degree reversal (Fig. 5F-G). Together, our work supports vector-like addition on collective electrotaxis.

We simplified the guidance cues at the edge of the cell sheet into two factors: default directional cues at the free edge and EF guidance. We can decompose the displacement of the cell sheet over 6 hours into components attributed to each guidance cue for the leading edge (EF + cues at free edge) and rear edge (EF – cues at free edge) of the cell sheet (Fig. S7A). With the known displacement of leading edge and rear edge of the cell sheet in cell experiments (Fig. 2B), we calculate the contribution of the free edge and EF guidance in those displacements, respectively. Compared with the displacement of the free edge we measured in cell experiment without EFs (Fig. 2F), there is a significant suppression effect of the free edge guidance when an EF is applied to cell sheets, and the higher EF, the stronger the effect (Fig.S7B).

### Propagation of migration speed and directionality of collective electrotaxis

EF-induced changes in migration speed and directionality propagate across the cell sheets, forming wave-like pattern, which are the results of integration of EF and free edge signals. In multi-cellular level, similar behaviors have also been demonstrated, like migration motility waves (Zaritsky et al., 2012; Zhang et al., 2017), traction force waves (Serrapicamal et al., 2012) in monolayer cell wound healing models, long-range force transmission during collective cell durotaxis (Sunyer et al., 2016), injury-induced Ca^2+^ waves (Klepeis et al., 2001), and ERK activation in epithelial cells initiated at the wound edge traveling as a wave to neighboring cells (Hiratsuka et al., 2015; Matsubayashi et al., 2004). Mathematical models have demonstrated similar behaviors across cell population (Bhowmik et al., 2016; Serrapicamal et al., 2012; Sunyer et al., 2016; Zhang et al., 2017). Our results from experiments and *in silico* modeling demonstrate the EF-induced wave-like propagation in both cell migration directionality and speed, initiated locally at the leading edge, and then globally (across the whole sheet). How those wave-like behaviors are initiated and regulated in collective electrotaxis will need further investigation. Possible mechanisms may include intracellular and intercellular signaling.

### Modeling suggests some mechanisms for collective electrotaxis

*In silico* model has suggested mechanisms for synergy and integration of different cues. Models of intracellular processes have been used in the chemotaxis field to understand how amplification, adaptation, and other processes allow single cells to transmit an external gradient into a gradient in signaling and cytoskeleton assembly (Masuzzo et al., 2016).

It is encouraging and satisfactory that with limited parameters, the *in silico* model produces the three distinct phases to wave propagation that matches experimental results: the initial phase (P1), ramping-up phase (P2), and saturation phase (P3).

Parameter test of our mathematical model indicates that the duration of P1 is mediated by a combination of cellular response time and strength of EF (Fig.S6A-B). In this phase, cells at the leading edge (i.e. cells originally migrated in the direction of EF) initiate the rising of directionality and speed in the field direction, which lead to more space as well as mechanical cues (e.g., pulling force) for the following cells. For P2, cells in the bulk of the cell sheet respond almost at the same time (Fig.6SA-B). For P3, our modeling results suggest that the characteristics of this phase is affected by cellular response time, strength of EF and also cellular motility state prior to EF stimulation. For example, it takes more time for a cell to follow EF guidance when its original migration direction was against the EF direction; similarly, cells in the rear edge of the cell sheet must integrate competing EF and default directional cues at the free edge through tug-of-war (Fig. 5E).

A delay in electrotaxis was found for cell sheets responding to the onset and the reversal of EF polarity (Fig.3C, 5A-D). Similar delays have been observed in responses of single cells when exposed to an EF or when the EF polarity is reversed (Sun et al., 2013). Prior research described this kind of delay as ‘memory,’ which has been observed in cells during chemotaxis. This memory effect plays an essential role in Dictyostelium cells in traveling waves of chemoattractant (Skoge et al., 2014; Wang et al., 2012). The directional sensing component activated Ras and its downstream targets, need time to adapt to new direction signals (Takeda et al., 2012). The delay effect develops the theory of the simple vector-like addition in collective decision making in direction (Fig. 5G). Vector addition accounts for integration of different cues in space, while the delay effect accounts for different cues in time. An EF of 50mV/mm dominates in the integration of electrical cue and default directional cues at the free edge in cell sheets of millimeters in size over ∼ 4-6 hours. The field strength and migratory responses are consistent with EFs measured at wounds and with the spatiotemporal features of cellular migration in wound healing.

### Limitations of study

It is worth noting that the results of sheets of human keratinocytes (HaCat cells) manifest major features of collective migration of keratinocyte sheet in wound healing. Those conclusions may need to be tested with other types of cells, especially primary human keratinocytes. However, we believe that the main conclusions and features are applicable to other types of cells, because we observed similar cell health status and motility in sheets of corneal epithelial cells and keratinocytes from human, bovine, mouse, and rat, in well-controlled culture systems (Zhao et al., 1996; Zhao et al., 2006). Those cell monolayers maintain cell-cell junction during collective migration and show no obvious cell death. In those models, as well as presented herein, leading edge cells are very motile and lead collective migration of whole sheet, similar to what have been found in various wound healing models (Eming et al., 2014; Gurtner et al., 2008; Martin, 1997; Pastar et al., 2014). The *in silico* model may benefit from including a Baysian framework to refine our estimation of unknown parameters, but we believe no significant qualitative addition would be likely by doing so, because our current model captures major key features manifested in experiments. The model may however benefit by considering molecular mechanisms of intracellular and intercellular signaling, thus, to gain insights to the directionality and speed waves induced by electric fields initiated at the leading edge.

## Author contributions

Y.Z., T.P., Z.P., F.L. and M.Z. designed the experiments, W.L, S.L., G.Z., Z.X., T.P., F.L. and M.Z. supervised the study. Y.Z. performed the cell experiment and analysis, G.X. and J.W. coded the mathematical model, R.M.L. programmed codes for analysis, Z.Z. designed and fabricated the chip. Z.Y., G.X., F.L., and M.Z. wrote the manuscript. All authors commented and/or edited the manuscript and figures.

## Acknowledgements

This work is supported by AFOSR MURI grant FA9550-16-1-0052, NEI R01EY019101, Core Grant (P-30 EY012576). This work was also supported by the National Science Foundation of China (Grant No. 51807142) and Hangzhou Normal University Research start-up Funds (No. 2019QDL031) to Y. Z., a Discovery Grant from the Natural Sciences and Engineering Research Council of Canada to F. L. (RGPIN-2014-04789). T.P. is supported by the Guangdong Program (2016ZT06D631), the Shenzhen Fundamental Research Program (JCYJ20170413164102261), Shenzhen Engineering Laboratory of Single-molecule Detection and Instrument Development (XMHT20190204002). MZ, YZ and FL would like to thank Alex Mogilner (NYU) for insightful discussion.

## Declaration of interests

The authors declare no competing interests.

### Supplementary Movie

**Video S1. Time-lapse phase contrast videos of directional collective migration of a square cell sheet in control group and 200 mV/mm EF group after peeling off a PDMS stencil, related to** **Figure 1**.

The timestamp is in minutes. Bar = 200 µm.

**Video S2. Time-lapse phase contrast videos of directional collective migration of a square cell sheet in 200 mV/mm group and cell migration trajectories of cells in regions of the cell sheet, related to** **Figure 2**.

The timestamp is in minutes. Bar = 200 µm.

**Video S3. Time-lapse video of directionality derived from PIV for control group and 200 mV/mm EF group, related to** **Figure 3**.

Color codes the cell migration directionality. The timestamp is in minutes. The still images of directionality shown in Fig. 3a, 3b are derived from Video S3.

**Video S4. Time-lapse video of speed derived from PIV for control group and 200 mV/mm EF group, related to** **Figure 3**.

Color codes the cell migration speed. The timestamp is in minutes. The snapshots of speed shown in Fig. 3a, 3b are derived from Video S4.

**Video S5. Simulation of the control group and 200 mV/mm EF group in the Particle-Based Compass model, related to** **Figure 4**.

Each ball indicates one cell. The dynamic color of the ball indicates the directionality. The snapshots of the computer simulation shown in Fig. 4e are derived from Video S5.

**Video S6. Time-lapse phase contrast video and video of directionality derived from PIV for 50 mV/mm EF group, related to** **Figure 5**.

Color codes the directionality. The timestamp is in minutes. Bar = 200 µm.

**Video S7. Time-lapse phase contrast video and video of and directionality derived from PIV for rEF group, related to** **Figure 5**.

Color codes the directionality. The timestamp is in minutes. Bar = 200 µm. 200 mV/mm EF stimulation started one hour after the start of image recording and the direction of EF stimulation reversed 3 hours later as indicated in the video. The duration of the video recording was 7 hours.

**Video S8. Time-lapse videos of flow vectors derived from PIV for rEF group, related to** **Figure 5**.

Arrows indicate the direction of cell migration. The length of the arrows indicates the speed of the cell migration.

## STAR ★METHODS

### KEY RESOURCE TABLE RESOURCE AVAILABILITY

#### Lead contact

Further information and requests for resources and reagents should be directed to and will be fulfilled by the lead contact, Min Zhao.

#### Materials availability

This study did not generate new unique reagents.

#### Data and code availability

- Raw microscopy data reported in this paper have been deposited at Mendeley Data and are publicly available as of the date of publication. The DOI is listed in the key resources table.
- All original code has been deposited at Mendeley Data and is publicly available as of the date of publication. DOIs are listed in the key resources table.
- Any additional information required to reanalyze the data reported in this paper is available from the lead contact upon request.

### EXPERIMENTAL MODEL AND SUBJECT DETAILS

#### Human skin keratinocyte cells

Human skin keratinocyte cells were cultured in Dulbecco’s modified Eagle’s medium supplemented with 10% FBS (Life Technologies) and 1% (v/v) penicillin/streptomycin (Life Technologies). Cells were seeded and maintained at 37℃ and 5% CO2 in air before experiments. During imaging, cells were cultured with medium supplemented with an additional 20mM/ml HEPES.

### METHOD DETAILS

#### Cell Patterning

Before seeding the cells, the stimulating region of the electrotaxis chamber was pre-coated with an FNC coating mix (an aqueous solution of fibronectin and other cell adhesion proteins) following the manufacture’s instruction to facilitate cell attachment. The area was rinsed with PBS twice and left to air dry. To seed cells in the stimulation region and control the shape and size of the cell sheet, PDMS stencils containing square microwells were utilized. The stencil was cut from a 250 μm thick sheet of silicone rubber (B&J Rubber Products, US) by a computer-controlled laser machine (VLS2.30, Engraver’s Network, US). The PDMS stencils were deposited on the surface such that the long axis of the stencil was parallel with the electrical stimulating channel, avoiding bubbles between the chamber and stencil. The PDMS stencil formed a reversible, watertight seal against the cell culture dish. For each electrotaxis chamber, two PDMS stencils were applied, one for the electrical stimulation group and the other for the control (without electrical stimulation). Cell suspensions were seeded with a density of 4×10^6^ cells/mL on top of the stencil and cells were allowed to attach for 30 min, then the cell suspension was gently replaced with fresh cell culture medium. After 14-16 h, cells formed a confluent monolayer. The stencils were carefully peeled off and the patterned cell sheets were left on the cell culture dish (Fig.S1A-C). Image recording started after the setup of the electrotaxis system, which took approximately 2 hours. Image recording ended within 24 hours after seeding cells, cell proliferation rate was less than 2.5%.

#### Electrotaxis system setup

The electrotaxis system was set up as previously published (Song et al., 2007) with minor modifications. The system is illustrated in Fig.S1D-F. Briefly, the electrotaxis chamber contains two cell culture regions, one for electrical stimulation and the other one for negative control. The cover glass was placed after the cell sheets had been formed on the dish. The electrotaxis system contains the electrotaxis chamber, salt bridges, saline, wires, and DC supply (Fig.S1D-E). The current density and electric potential of the electrotaxis chamber were simulated by COMSOL Multiphysics, EFs are relatively evenly distributed in the major EF stimulation area (Fig. S1F).

#### Time-lapse image recording

Cell migration was monitored with an inverted microscope (Carl Zeiss, Oberkochen, Germany) equipped with an XY motorized stage, time-lapse imaging software (Metamorph NX; Molecular Device, Sunnyvale, USA), and a Carl Zeiss incubation system. The microscope system was able to capture images of multiple locations. For recording cell sheets’ collective migration, 9 to 12 regions were recorded for each cell sheet allowing about 20% percent overlap for each region. Images were then stitched together using ImageJ software from the National Institutes of Health. All experiments were conducted using 10 × phase contrast objectives. Images are taken at 5 min intervals.

#### Microfluidic chip experiment setup and analysis

Human keratinocyte cells were cultured routinely to form a confluent monolayer in Petri dishes. A multi-electrodes microfluidic chip was fabricated to generate patterned electrical fields. A vacuum-assisted assembly method was utilized to allow direct attachment of our device to the existing cell monolayer, as was described before (Zhao et al., 2014). COMSOL Multiphysics was employed to simulate the electrical field lines in chips in the 2D mode of EF in conductive material, the conductivity of cell culture medium was set to 1.3 S/m. Cell migration was monitored by phase contrast microscopy on an inverted microscope equipped with a motorized stage, time-lapse imaging software, and Carl Zeiss incubation system. PIV (Particle image velocimetry) analysis and Line Integral Convolution (LIC) assay were also adopted to quantitate and illustrate the temporal and spatial dynamics of cell collective migration.

#### Math model

The PBC model for the electrotaxis assay was modified from our previous model that was used to simulate a wound healing assay (Zhang et al., 2017). The parameter values were chosen because that they are in biologically realistic ranges based on our experimental data, and refined by testing of the model to make improved match to the cell experiment result. For instant, in the experiments, the cell cluster is about 1mm by 1mm in size and each cell is about 20 µm in diameter, so there are about 2500 cells in the cell cluster. Accordingly, there are also 2500 cells in the model, the layout is a square shape with 50 rows and 50 columns. Based on these experimental data, we configured total 2500 cells in a 50 by 50 matrix layout in the model. As another example, based on cell diameter of about 20 µm and cell speed of about 20 µm to 30 µm per hour observed in the experiments (which is about 1.5 times of cell diameter), we in the model set the cell diameter to be 0.1 and cell speed to be 0.15 per hour to be consistent. The EF speed in low EF condition is lower than that of high EF, and it takes longer time for the cells to reach maximum EF speed in low EF as well. While parameters such as cell-cell interaction related parameters, delay effect related parameters, the value are determined based on tests and refined through comparison with experiment results. There is no relative position changes during simulation as it was stated in cell experiment. In our cell experiment, over 90% cells in the cell sheet are surrounded by eight neighbors, to be consistent with biological data, we set eight surround sectors for our modelling and the cell’s direction of motion is also determined by the nearest neighbor cells divided into eight sectors. If there is no neighbor cell in one or more sectors, the corresponding sectors are marked as empty sectors. The cell-cell interaction is either attraction or repulsion depending on the distance between the cells. Empty sectors always have the strongest attraction force. Details of the cell-cell interaction strategy with its nearest neighbors can also be found in our previous paper (Zhang et al., 2017). When EF was applied in this electrotaxis assay, we directly added an EF-induced displacement (EFspeed) in addition to the displacement caused by cell-cell interactions (CellSpeed):]

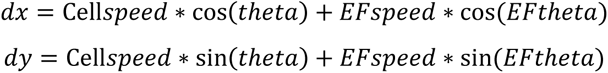

Theta is the direction determined by cell-cell interactions, EFspeed is the magnitude of the EF effect on the cell displacement, and EFtheta is the direction of the EF anode added with random noise. In the model, a delay coefficient variable preWeight is used to determine how much percentage of previous moving speed is kept at the current time step, the following formula shows how to compute cell’s total speed at current time step t:

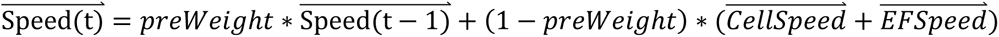

The preWeight parameter is tuned and 80% is selected for preWeight to align with experimental results. To simulate the additional delay effect for cellular response to the EF guidance, EF-induced displacement (EFspeed) in the model increase linearly from zero to the maximum value of the EFSpeed after turning on EF stimulation and decrease also linearly after switching off EF stimulation. The cellular response time to EF guidance (20 min for high EF and 120 min for low EF) is determined based on data of single cell experiments. There are 220 time steps in the model, the cell array is allowed to have 10 time steps to warm up to be more close to the state of cell sheet in reality. The following 210 time steps in the model stand for the 7 hour in cell experiments. MATLAB (MathWorks) was used as the programming environment to implement the simulations. The parameter table of the modeling is provided in the Supplementary Material (Table S1). All original code has been deposited at Mendeley Data.

### QUANTIFICATION AND STATISTICAL ANALYSIS

#### Individual cell tracking

Individual cells were tracked using ImageJ’s MTrackJ plugin (Schindelin et al., 2012). To analyze region dependent cell migration in the cell sheet, we defined cells in the leading region, rear region, and lateral region (right) as the 10 rows of cells away from the edge. The position of a cell was defined by its nucleus. For each region, 20 unconnected cells were tracked at 10 minutes frame intervals. The directionality was used to quantify the directedness of cell migration, which was defined as the cosine of the angle between the cell motion and the y axis. In our experiment, the electrical field was applied along the y axis directed down the image (cathode at the top and anode at the bottom). A cell migrating directly toward anode would have cosine = 1 and sine = 0. The cell migration rate was quantified as the accumulated migration displacement along the y axis.

For cell relative position tracking, we selected 20 cells that are not neighbors in each region (front, back, central, left and right). Neighboring cells of those 20 cells were identified manually, tracked throughout the video. If the neighbor cells maintained their relative position to the selected cells, we concluded that the cells did not change their relative position. Otherwise, if the neighbor cells did not maintain their relative position with the selected cells, we conclude the cells changed its relative position. The percent of cells that change their relative position divided by the total number of cells tracked is thus calculated.

For the computational model, we exported cell coordinates (which indicate the cells’ current positions) with cell indexes (which are correlated to the cells’ initial positions in the cell sheet) at selected time steps of the simulations. Then we sorted the X coordinates for each row of cells in numerical order for each selected time points and checked if their cell indexes after sorting remained the same order. If the order matches, then we conclude no relative position change in the X direction. The similar method was applied to the Y coordinates and index of each column of cells to evaluate the relative position change in the Y direction.

#### Particle image velocimetry

Heatmaps and kymographs of directionality and speed were generated using a custom MATLAB code based on MatPIV1.6.1, a freeware distributed under the terms of the GNU general public license. The MATLAB code has been previously described in detail (Sveen, 2004). Kymographs were used to quantify and visualize spatiotemporal dynamics of directionality and speed from the PIV measurements. For each data matrix from the PIV analysis, we computed the average value for each column or row parallel to the edge and then derived a one-dimensional segment for each time point.

#### Contour and centroid tracking of cell sheet

The contour of the cell sheets was segmented by custom MATLAB code that uses sobel filtering to find edges in the images. Small, segmented regions such as those from individual scattered cells were excluded. The centroid of the cell sheet was computed using MATLAB’s regionprops function.

#### Statistical analysis

Data analyses, graphs, and statistical calculations were performed using Excel (Microsoft) and MATLAB. Data are presented as mean ± standard error of the mean (SEM). Differences between conditions were compared using an unpaired Student’s *t*- test. The criterion for statistical significance was *p*<0.05.

