## Supplemental material for "Propagation dynamics of electrotactic motility in large epithelial cell sheets"

**Supplementary Materials**

**Table S1. Parameters in the PBC model, related to STAR Methods.**

| Symbols | Descriptions | Values (unit) |
|---|---|---|
| cell_num | Total number of simulated cells | 2500 |
| cell_speed | Cell speed | 0.005 |
| angle | Cell migration angle | It is calculated based on cell migration distance along x and y axes. |
| R_R ($D_0$) | Repulsive threshold distance | When cell-cell distance is less than $D_0$, it is repulsive interaction. $D_0$= 0.1； For reverse EF condition, $D_0$ is adjusted to 0.13 after EF reversion to compensate the cell-cell distance increase in the 1st half experiment. |
| attractiveTH ($D_{max}$) | Attractive threshold distance, threshold of the free edge effect | When cell-cell distance is between $D_0$ and $D_{max}$, there is an attractive interaction increases with distance; when cell-cell distance is equal to or greater than $D_{max}$ (or cell at free edge), the attractive interaction reaches its maximum. $D_{max}$ is set to 0.3 in the simulation. |
| preWeight | Weight of cell speed at the previous time step | 80% |
| high_ef_speed | The magnitude of EF speed for the 200mv simulation | 0.005 |
| low_ef_speed | The magnitude of EFspeed for the 50mv simulation | 0.0033 |
| tmax | Total time steps | 220 |
| efangle | The direction of ef speed | 0.5*pi or -0.5*pi (reversed EF) |
| dx | A vector to keep all cells' x coordinates | vector, size is 1 by cell_num |
| dy | A vector to keep all cells' y coordinates | vector, size is 1 by cell_num |
| totalspeed | Cell's total speed, if there is no ef signal applied, totalspeed is equal to cell speed. If ef signal is applied, totalspeed is the vector addition of cell speed and ef speed | totalspeed = cell_speed + EF_speed |
| vx | A matrix to keep all cells' x coordinates at all time steps | Matrix, the shape is cell_num by tmax |
| vy | A matrix to keep all cells' y coordinates at all time steps | Matrix, the shape is cell_num by tmax |

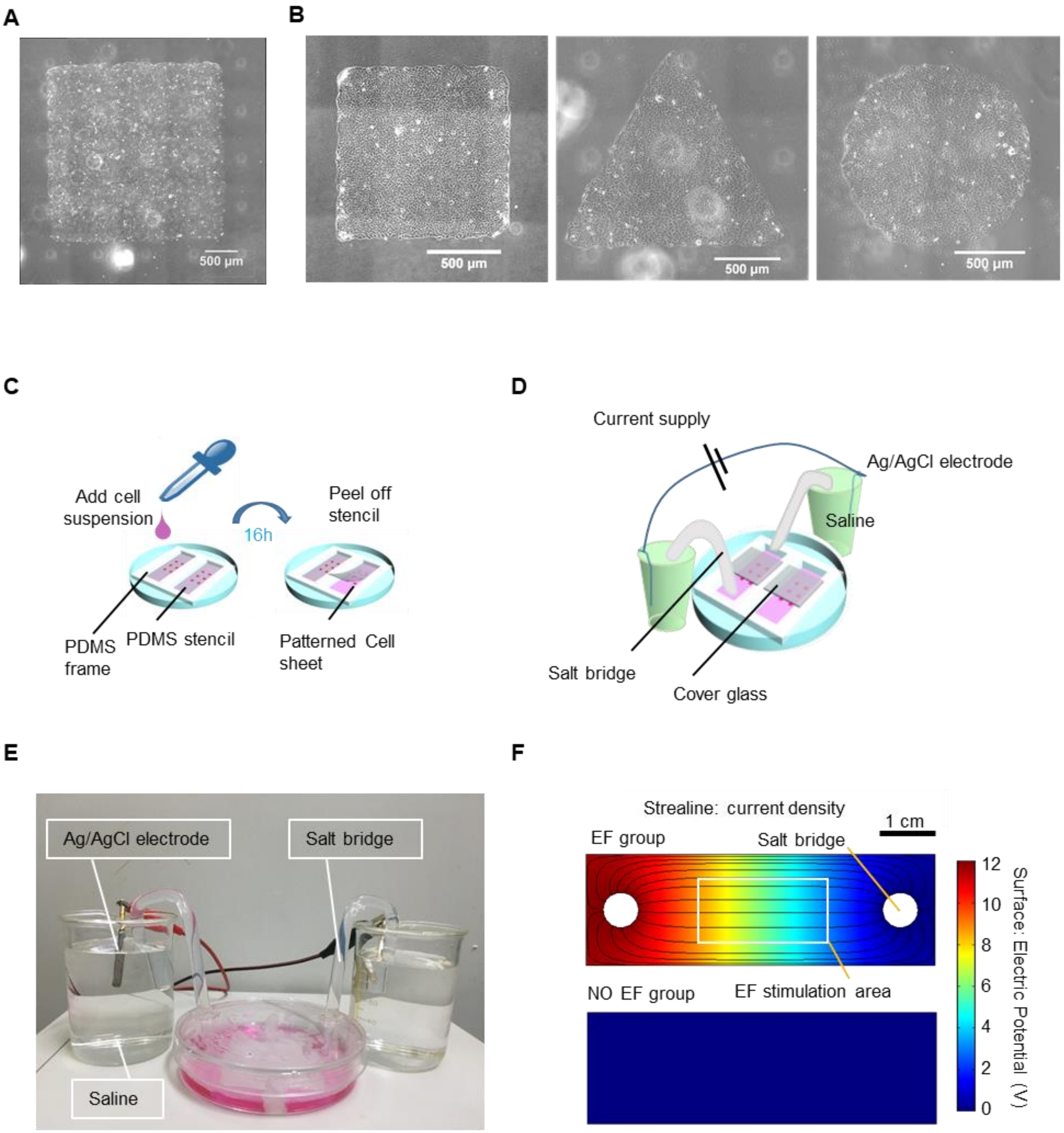


**Figure S1. Engineering of cell sheets and experiment setup, related to Figure 1.**
**A-B** Cell sheets of defined size and shapes engineered using PDMS stencil. Scale bar = 500 µm. **C-E** cell plating workflow and electrotaxis experiment setup. **F** Electric potential and current density of electrotaxis chamber simulated by COMOSL Multiphysics. Scale bar = 1 cm.

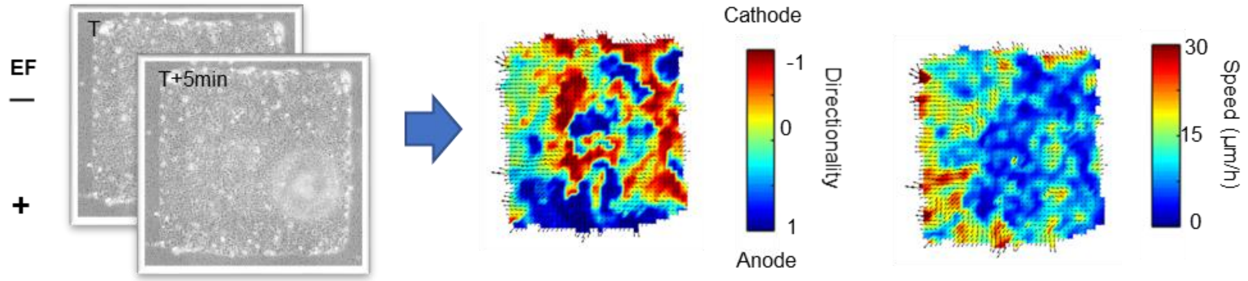


**Figure S2. Directionality and speed of movement of the cell sheet shown as heatmaps from PIV analysis from two adjacent images with a time interval of 5 min, related to Figure 3.**

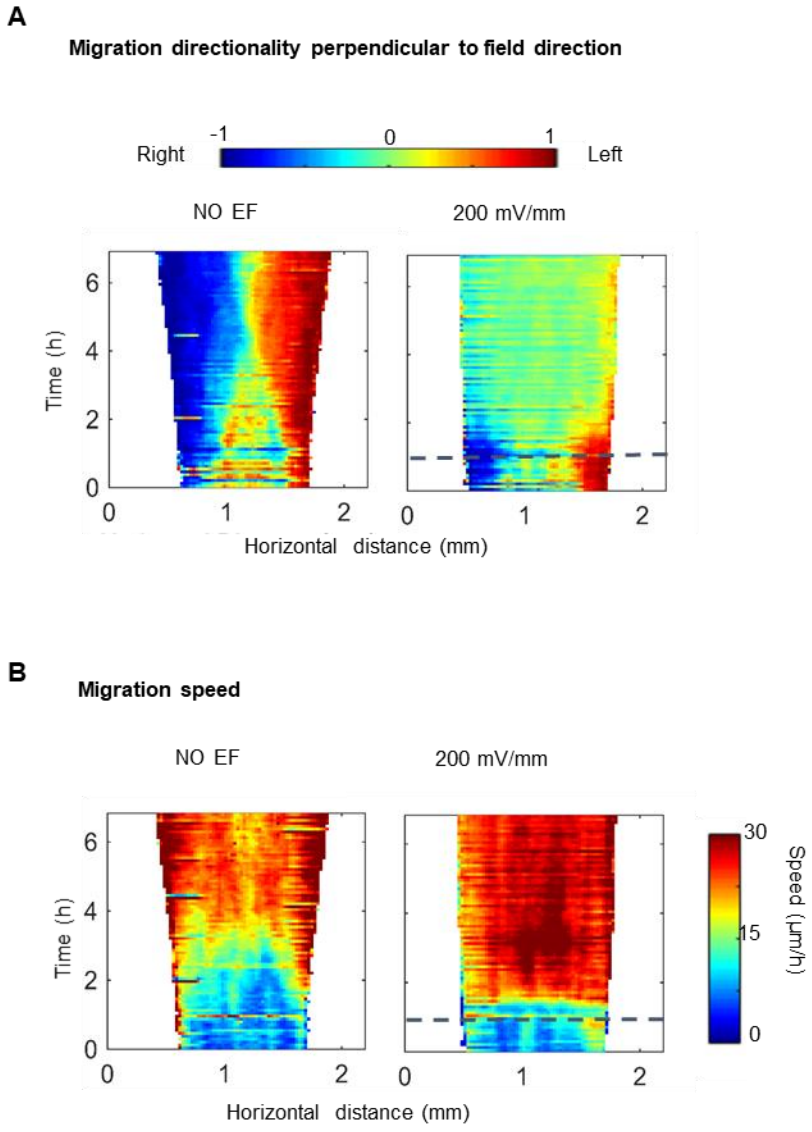


**Figure S3. EFs suppress the guidance effect of free-edge perpendicular to the field line (vertical) and abolish the free-edge induced directionality wave, related to Figure 3.**

Kymographs of migration directionality (**A**) and speed (**B**) as a function of time (y axis), respectively. Kymographs in NO EF group show that speed dynamic induced by free edge propagating from edge to the center of the cell sheet. An EF significantly decreases the directionality along the x axis. Dashed lines indicate the onset of the field. The experiment duration is 7 hours. Kymographs are from one of three independent experiments with the same pattern.

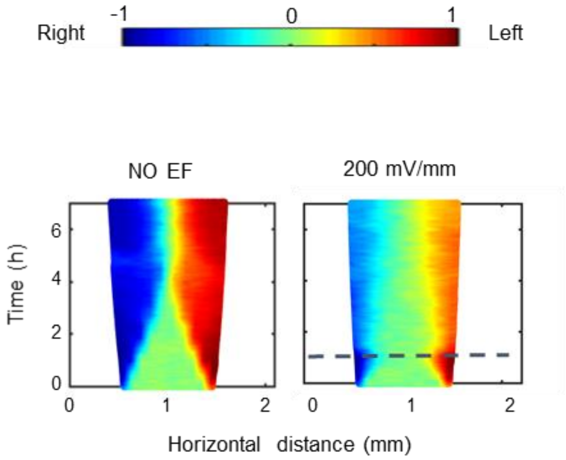


**Figure S4. The PBC model replicates the suppression effect of EF on cell sheet expansion in the direction perpendicular to the field line, related to Figure 4.**
*In silico* modeling show migration directionality in kymographs. In the NO EF group, increases in migration directionality initiate from the free edge and propagate from the edge to the center of the cell sheet. An EF suppresses and almost abolishes the increase in directionality along the x axis (compare with Fig. S3A). Dashed lines indicate the onset of the field. The experiment duration is 7 hours.

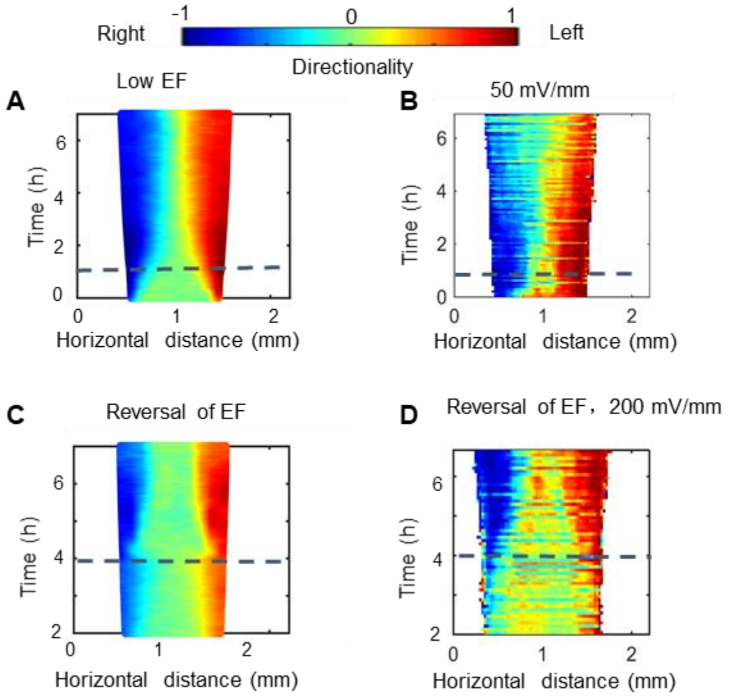


**Figure S5. Spatiotemporal dynamics of collective electrotaxis in the PBC model, related to Figure 5.**

**A-D** *In silico* results (**A**, **C**) faithfully predict the dynamics of migration directionality in 50 mV/mm EF group and reversal EF group (**B**, **D**). Grey dash lines indicate the onset and reversal of the field. Kymographs presented are from a representative experiment and were confirmed in three independent experiments.

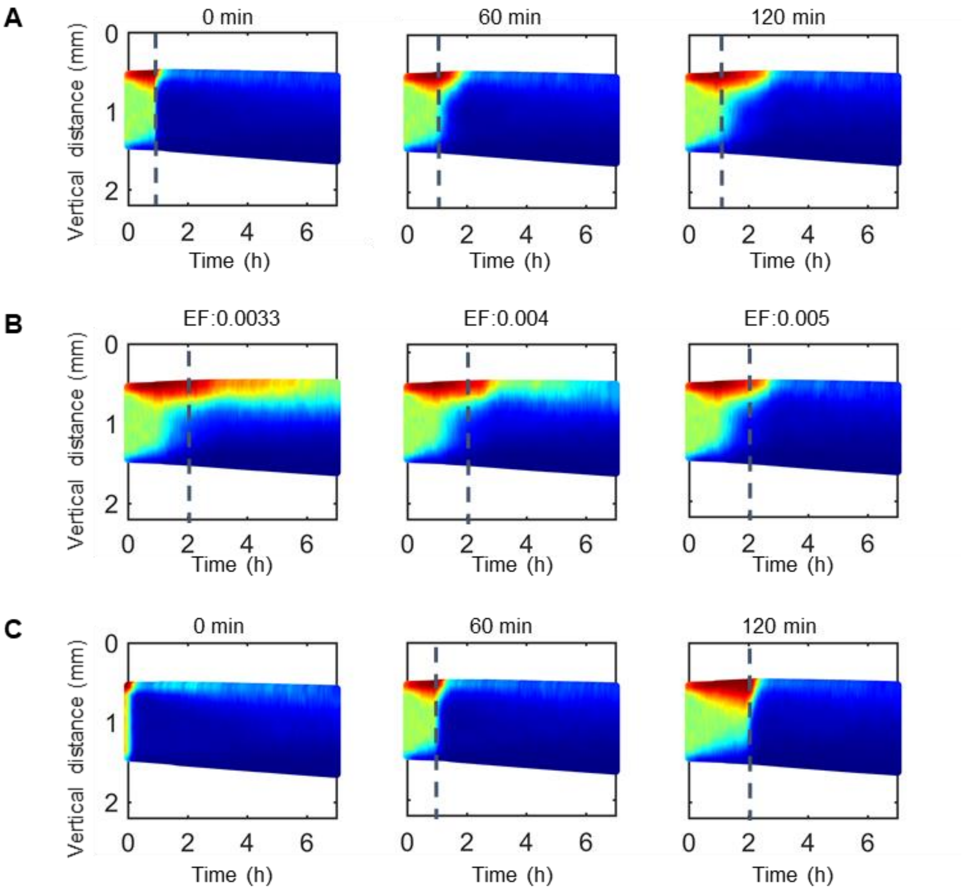


**Figure S6. *In silico* model implies the factors that affect the three-phase propagation dynamics of collective migration in EFs, related to figure 5.**

*In silico* results predict the different cellular response time to EF guidance (**A**), the strength of EF (**B**), motility state (**C**) affect the propagation dynamics of collective electrotaxis. (**A**) Cellular response time for EFs is set to 0 min, 60 min, and 120 min, in which the value of the EF increases from 0 to 0.005 linearly in the model. (**B**) Cellular response time for EF is set to 2 hours and strength of EF is set to 0.0033, 0.004, and 0.005, respectively. (**C**) We simulate the different motility state before turning on EF by adjusting the time point we start EF stimulation. Cellular response time is set to be 20 min and strength of EF is set to 0.005 in the model. Grey dash lines indicate the onset of the field. Kymographs presented are representative one confirmed in three independent experiments.

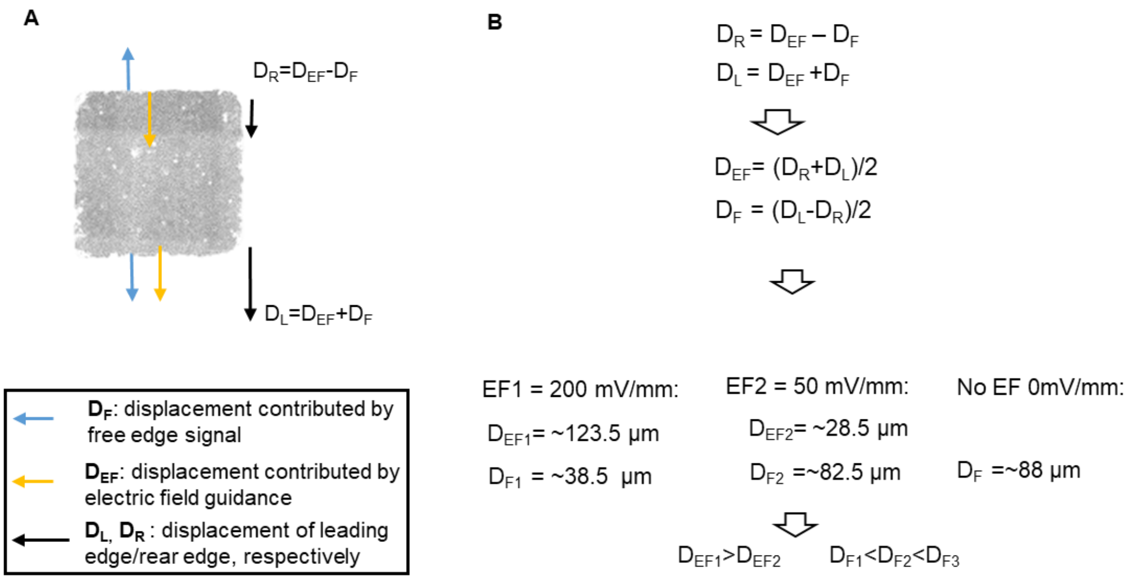


**Figure S7. Effect of suppression of the electric fields over free edge guidance, related to Figure 2.** **A** The guidance mechanisms at the edge of cell sheet are simplified as two cues: free edge and EFs. Displacement of the edges is decomposed over 6 hours into components attributed to each guidance cue for the leading edge ($D_L$=$D_{EF}$ + $D_F$) and rear edge ($D_R$=$D_{EF}$ - $D_F$). **B** Based on displacement of the leading edge (~162 µm) and rear edge (~85 µm) (experimental data from Fig. 2B, of cell sheets in 200 mV/mm EF), contributions were calculated for the free edge (~38.5 µm) and EF guidance (~123.5 µm) in those displacements, respectively. Using the known displacement of the leading edge (~111 µm) and rear edge (~-54 µm) of the cell sheet in 50 mV/mm EF (Fig. 5E), we do the same calculation for the 50 mV/mm EF group, and we find the contribution of free edge (~82.5 µm) and EF guidance (~28.5 µm) in those displacements, respectively. The displacement of free edge we measured in cell experiment of NO EF group is ~88 µm (Fig. 2F).
